# Learning stabilizes temporal activity but not neuronal selectivity in prefrontal cortex

**DOI:** 10.64898/2026.07.23.740357

**Authors:** Yu-Yang Huang(黄宇阳), Leonie S. Mehrke, Tobias W. Bernklau, Laura Busse, Simon N. Jacob

## Abstract

Intelligent behavior requires neural representations to change with new demands while preserving learned structure. The prefrontal cortex is central to this ability, but the mechanisms are unclear. Here, we tracked medial prefrontal neurons for months as mice learned an association task with successive rule switches. Learning progressively stabilized when individual neurons were active during a trial, but not what task variables they responded to. Neurons repeatedly gained, lost, or changed selectivity even after their activity profile had stabilized. We developed Sparse Tensor Component Analysis to show that, rather than reflecting random drift, dynamic single-neuron selectivity arose through rule-dependent recombination of a fixed set of task representations. Thus, learning established a stable temporal scaffold in the prefrontal cortex within which neurons participated flexibly in different representations.

## Introduction

A hallmark of intelligence, in biological and artificial systems alike, is the ability to retain what has been learned while continuing to learn something new. The prefrontal cortex (PFC) is central to this flexibility: it forms task representations during learning and updates them as the relations among cues, actions and outcomes change (*1–8*). Selectivity for the task variables that define a to-be-learned association develops in prefrontal neurons and strengthens over training (*9, 10*), so that neurons respond more distinctly across task conditions at expert performance (*11–17*).

Longitudinal recordings in the hippocampus and several neocortical regions show that neuronal responses associated with specific task events change across days even when behavior remains stable, a phenomenon termed representational drift (*18–23*). This continual reconfiguration has been proposed to support learning and flexibility by keeping alternative neuronal configurations available for new associations (*24–26*). Paradoxically, drift is not a prominent feature of the PFC. Longitudinal studies instead found temporal activity patterns that persist across days and even across changes in task contingencies, i.e., rules (*27–29*). How can a region central to learning and adaptation show such stable activity? More generally, what property of a prefrontal neuron’s response changes across learning, and what does learning stabilize?

Answering this question requires distinguishing two properties of a neuronal response that are typically conflated and treated as one. Activity is a neuron’s firing-rate profile over a trial: how strongly, and when, it responds. Selectivity is the difference between its responses across conditions of the same task variable: what cue, action, or outcome it distinguishes. The two need not change together. A neuron can retain the same overall activity profile while changing which condition evokes the stronger response; conversely, it can preserve a condition preference while shifting when it is active. Distinguishing the timing from the content (the “when” from the “what”) in cognitive coding is difficult in many behavioral tasks because task variables change as the trial unfolds. For example, an animal’s position in a maze covaries with elapsed time, so a shift in when a neuron responds can look like a change in what it represents (*18, 21, 24, 27, 28*). Reports of stable or drifting responses therefore do not, by themselves, reveal whether learning acts on neuronal activity, selectivity, or both (*30*).

Separating activity from selectivity leads to two accounts of prefrontal learning. In the first, activity-selectivity coupling, learning binds selectivity peaks in time to activity peaks. When a rule switch invalidates the learned association, activity and selectivity should be disrupted together before a new code forms (H_0_, **Fig. 1A**). In the second, activity-selectivity uncoupling, learning stabilizes when neurons are active, creating a temporal scaffold for the evolving trial, while selectivity can be reassigned as task demands change. A rule switch should then alter what individual neurons distinguish without erasing when they respond (H_1_, **Fig. 1B**). This second account would allow the PFC to preserve the structure of a task while flexibly changing the associations implemented within it. Distinguishing these accounts thus requires following the same neurons throughout learning and, importantly, across changes in task rules, while examining activity and selectivity separately.

**Fig. 1.**
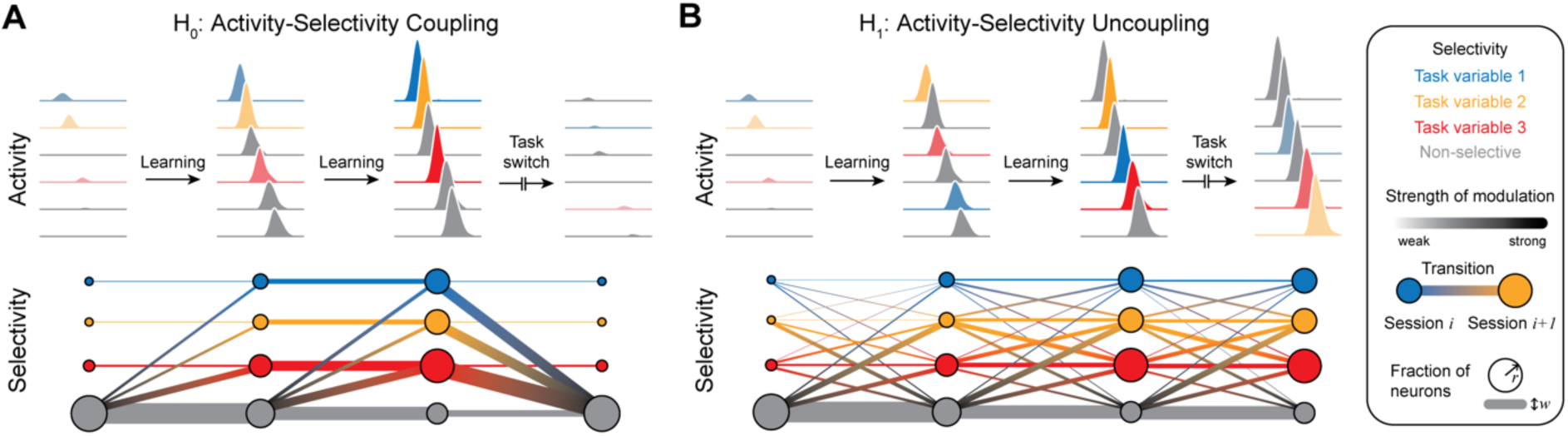
Competing accounts of learning. (**A**) Activity-selectivity coupling (H_0_). Individual neurons display peaks of activity that are localized in time and tile the trial. Activity increases during learning, but retains its peak timing. The peak of activity temporally coincides with the peak of selectivity. Selectivity increases during learning. Neurons retain their selectivity and express it persistently. After a switch of task contingencies, both activity and selectivity collapse, and neurons return to the non-selective state. (**B**) Activity-selectivity uncoupling (H_1_). Individual neurons display temporally localized peaks of activity as in H_0_. Selectivity increases during learning. However, neurons switch their selectivity from session to session and only express it transiently. After task switches, activity is retained, and changes of selectivity reflect the new task demands.

We tested these conceptual extremes by tracking the same medial PFC (mPFC) neurons for months as mice learned three successive cue-action-outcome associations (*31*). The rule switches changed which cue predicted each action and outcome but preserved the sequence and timing of events within a trial, allowing changes in task variable selectivity to be separated from changes in temporal activity. Learning progressively stabilized when individual neurons were active, and these temporal profiles persisted across rule switches. By contrast, the selectivity of individual neurons did not stabilize, neither within nor across rules (in line with H_1_, **Fig. 1B**). To identify the source of this transient selectivity, we developed Sparse Tensor Component Analysis (STCA), a method that decomposes population activity into a set of recurring task representations and separates, for each one, which neurons contribute from how strongly they are engaged in a given session. Single-neuron selectivity changed across sessions not because neurons changed their functional identity, but because the PFC recombined task representations as the rule changed, recruiting a given neuron into some of the representations in one session and releasing it in another. This arrangement allowed prefrontal neurons to retain the structure of a learned task through their stable activity profile, while encoding new associations through changes of their selectivity. Thus, the network could adapt without rebuilding what it had already learned. Together, our findings extend beyond the rodent prefrontal cortex and demonstrate how a cortical neural circuit bears on a challenge common to any adaptive system evolved for intelligent behavior.

### Rule-switching task and longitudinal neuronal recordings

To investigate how learning and changing task demands shape single-neuron and population-level representations in mPFC, we trained head-fixed mice (*N* = 12) to perform a cue-action-outcome association task with implicit, uncued rule switches. In each trial, an auditory stimulus (cue) of either high or low frequency noise was presented from either the left or right side (**Fig. 2A**). Following the cue, two spouts advanced toward the mouse, and the animal had to lick the correct spout to receive a water reward. Whether a trial was correct or false depended on the current task rule. The trial structure, i.e., the sequence and timing of cue, response and outcome, remained unchanged for all rules. The animals acquired three task rules sequentially through trial-and-error (**Fig. 2B**). The animals were first trained to follow cue location (left or right; R1), followed by cue frequency (low or high; R2), and finally cue frequency with reversed cue-action mapping (R3). The entire training duration spanned up to three months (one session per day) (**Fig. 2C**). Sessions were categorized by performance into novice, intermediate and expert learning stages (below 60 %, 60 to 70 % and above 70 % accuracy, respectively).

**Fig. 2.**
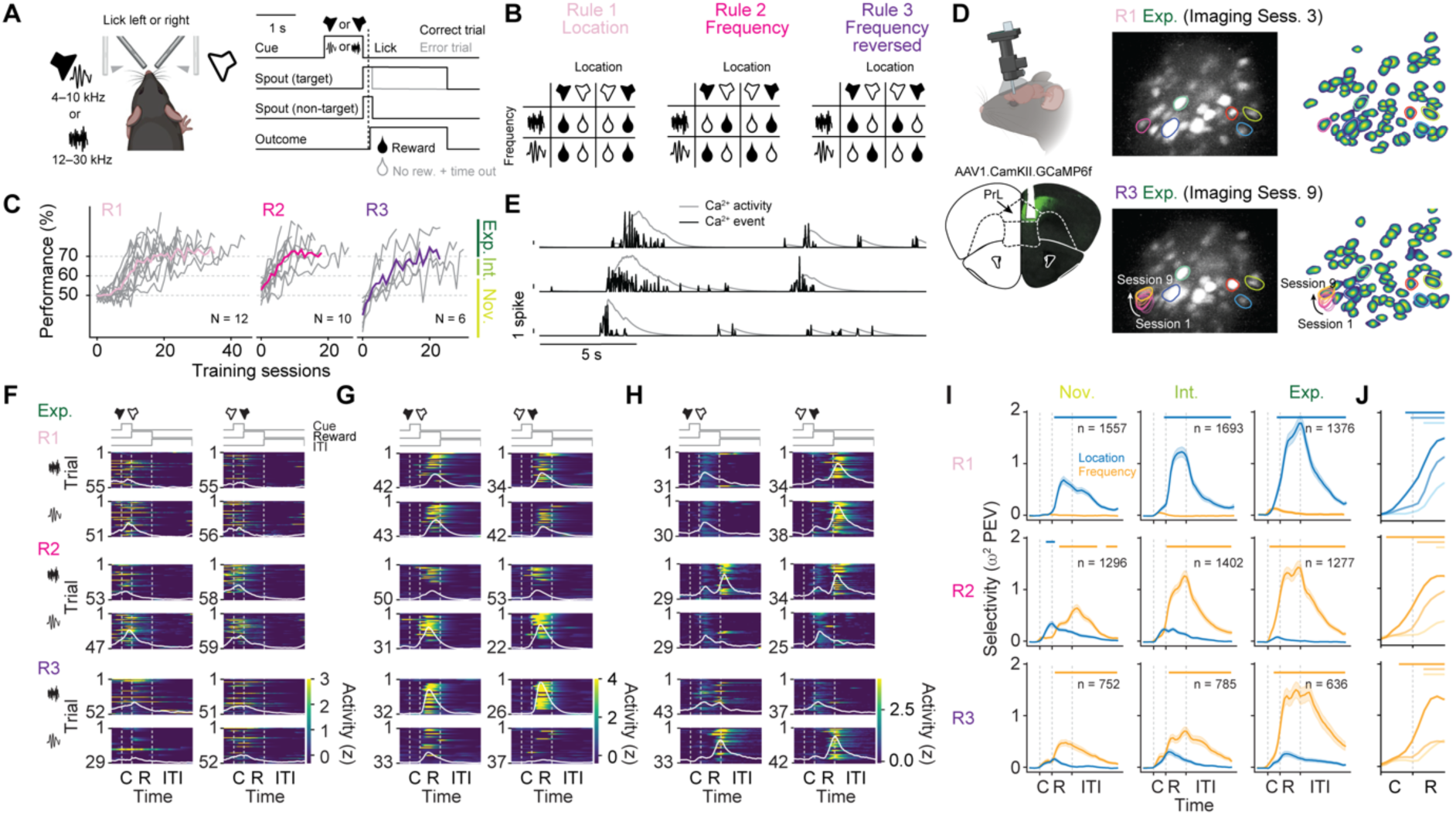
Behavioral task and longitudinal neuronal recordings. (**A** and **B**) Cue-action-outcome association task. Mice responded to an auditory stimulus (cue) by licking either the left or right spout for reward. The animals acquired three task rules sequentially (R1, location; R2, frequency; R3, frequency with reversed cue-action mapping) through trial-and-error. (**C**) Single-animal learning curves and group means. Cut-offs for novice, intermediate and expert learning stages are marked. (**D**) Miniscope calcium imaging over mPFC (prelimbic cortex, PrL). Fields of view (maximal intensity projection) and spatially registered neurons from the R1 and R3 expert sessions (top and bottom, respectively). Colored contours mark example neurons. One neuron is shown registered across all nine imaging sessions. (**E**) Calcium traces and deconvolved calcium events from three example neurons. (**F** to **H**) Activity of three example neurons, z-scored to the entire session, in correct trials of each rule’s expert session, split by condition. Solid and dashed lines mark trial-averaged activity and trial epochs, respectively. (**I**) Selectivity for cue location and cue frequency (expressed as ω² percent explained variance, PEV), averaged across all neurons, split by learning stages and rules. (**J**) Magnification of cue-epoch selectivity. Darker shades indicate later learning stages. Horizontal bars mark time points with significant differences between location and frequency selectivity (Wilcoxon signed-rank test, Bonferroni-corrected, family-wise error rate < 0.05). Error bands, SEM across neurons. C, cue epoch; R, response epoch; ITI, inter-trial interval.

We recorded the activity of a total of 3,556 mPFC excitatory projection neurons (GCaMP6f expressed under the CamKII promoter) across learning using chronic microendoscopic single-photon calcium imaging (**Fig. 2D**). For each task rule, we acquired imaging data in all three learning stages. We retained 2,604 units that were imaged in more than one session for further analysis, as these showed significantly increased signal-to-noise ratio and spatial footprint consistency compared to single-session units (**fig. S1**). For STCA, calcium activity was deconvolved to estimate putative spike events (*32, 33*) (**Fig. 2E**). Each neuron’s activity was normalized by z-scoring, using its session-specific mean and standard deviation. 88 % of neurons (2,299/2,604) exhibited task-related activity, i.e., changes in activity across trial epochs (pre-cue, cue presentation, response, inter-trial interval; one-way ANOVA, *p* < 0.05 in at least one session). Responses of individual neurons varied with cue location and cue frequency and changed with rule switches, indicating they tracked task contingencies and the animal’s behavior (Fig. 2F to **H**).

To quantify task variable selectivity, we computed effect sizes (*ω*^2^, percent explained variance) for each neuron using a two-way ANOVA, with cue location and frequency as main factors. Unless stated otherwise, we included correct and error trials in our analyses to comprehensively capture the learning process. In R1, population-averaged selectivity for cue location gradually increased with learning, while selectivity for cue frequency remained low (**Fig. 2I**, top). After the switch to R2, selectivity for cue location dropped sharply and continued to decline, whereas selectivity for cue frequency rose steadily (**Fig. 2I**, middle). Because the presented cues were identical across rules, this suggests that the observed selectivity reflected emerging decision-related processes rather than pure sensory encoding. Similar selectivity time courses were observed after the switch to R3 (**Fig. 2I**, bottom). Notably, with learning, the onset of task-relevant selectivity shifted forward in time from the response epoch to the cue, indicating that the animals had successfully acquired the cue-action-outcome association (**Fig. 2J**). Together, these results show that prefrontal population-level selectivity dynamically evolved with learning and reflected changing task demands.

### Stable population representations with reoriented projections after rule switches

We next investigated how the mPFC transformed sensory cue information into behavioral choices and how learning shaped this transition. Of the 12 implanted mice, six completed all three rules, yielding nine imaging sessions per animal and 54 in total. Two animals contributed less than 10 neurons that were recorded across all sessions and were therefore not included in further analyses. In the remaining four mice (performance from novice to expert, R1: 51 ± 1%, 62 ± 2%, 80 ± 1%; R2: 53 ± 3%, 63 ± 4%, 77 ± 3%; R3: 37 ± 3%, 64 ± 6%, 78 ± 2%), we recorded 170 neurons across all nine sessions. Of these, 149 neurons were task-related (one-way ANOVA across trial epochs, *p* < 0.05 in at least one session). We performed cross-temporal decoding, training binary support vector machine (SVM) classifiers to predict cue location (left or right), cue frequency (high or low) and behavioral choice (lick left or right) on a trial-by-trial basis (*34*). Across animals, decoding accuracy for all task variables increased with learning (**Fig. 3A**). During the cue epoch, decoding of the task-irrelevant sensory dimension (i.e., cue location in R2 and R3, and cue frequency in R1) also exceeded chance level, mirroring the brief rise in population selectivity above baseline for both relevant and irrelevant sensory dimensions during cue presentation (**Fig. 2I**). This effect persisted after error trials were excluded (**fig. S2A**). In the response epoch, decoding of the task-relevant sensory dimension closely tracked choice decoding, indicating that the corresponding sensory and choice coding axes were aligned.

**Fig. 3.**
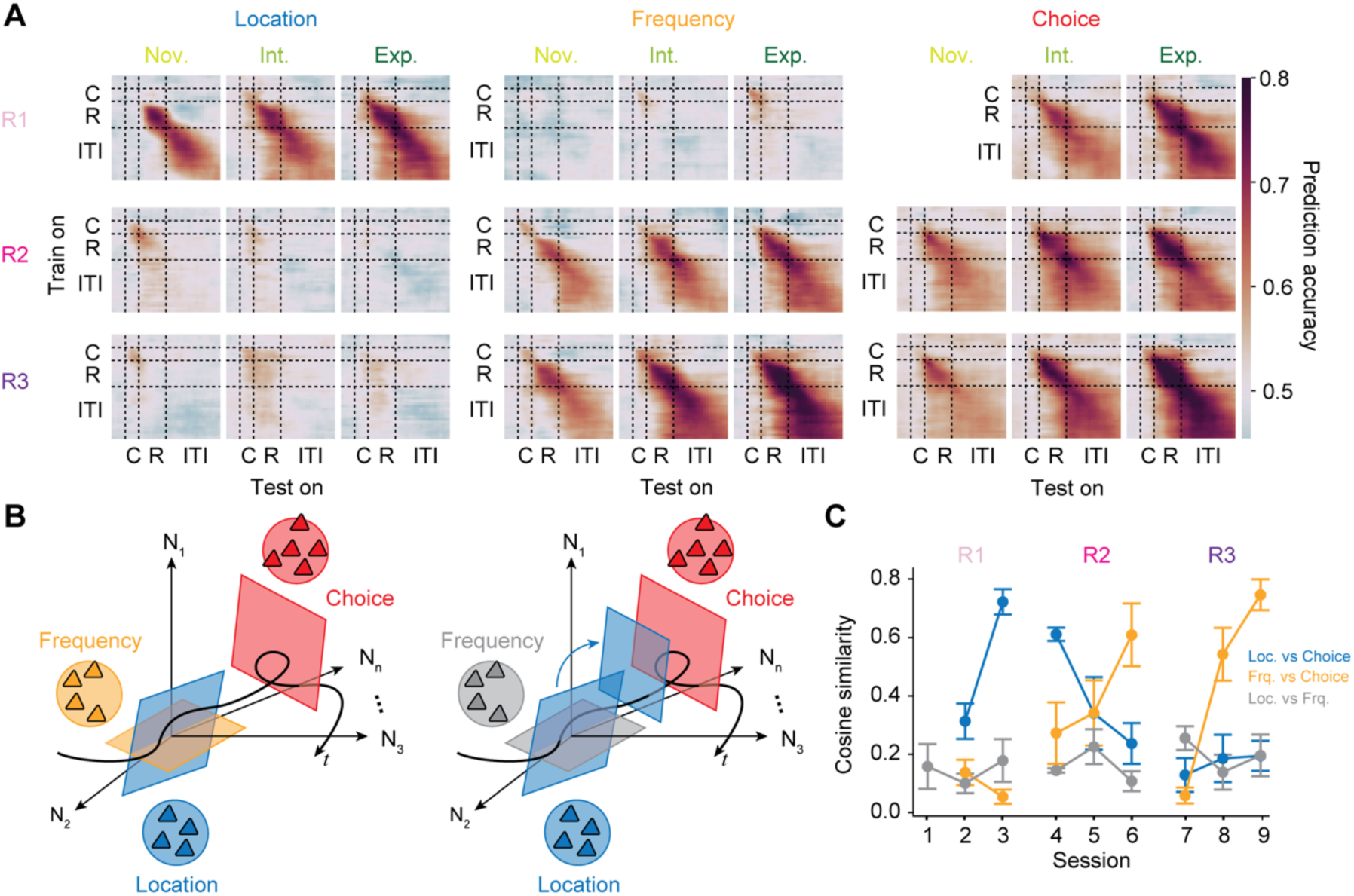
Transition from sensory cue to behavioral choice. (**A**) Prediction accuracy of linear binary SVM classifiers decoding cue location, cue frequency and behavioral choice, trained at each trial timepoint and tested at all timepoints within a session (*N* = 4 animals, *n* = 149 task-related neurons; correct and error trials). Choice decoding is omitted for R1 novice, because strong response biases resulted in too few trials of the non-preferred choice for conditions to be balanced. (**B**) Schematic of within-trial population dynamics in the expert stage (example for R1). Task variables are represented in distinct subspaces (left); as the trial progresses, the task-relevant cue subspace aligns with the choice subspace (right). (**C**) Decoder hyperplane alignment, defined by the cosine similarity between hyperplanes during the second half of the response epoch within each session, plotted across sessions for each rule. Error bars, SEM across animals.

These results suggested that mPFC neurons encode multiple sensory subspaces early in the trial, regardless of their task relevance, and that the task-relevant subspace is then selectively projected into the choice subspace as learning progresses (**Fig. 3B**). We examined how these representations evolved across learning using cross-session decoding (**fig. S2B** and **C**). Stability across sessions would argue for successful generalization (*35*). Choice decoders generalized well across sessions. For instance, a decoder trained in R1 predicted choices in R2 (**fig. S2B**), and a decoder trained in R2 generalized to both R1 and R3 (**fig. S2C**), suggesting that choice representations were stable and shared across rules despite contextual changes. By contrast, cue location decoding in R1 did not generalize after rule switch to R2 (**fig. S2B**). Notably, cue frequency and choice decoders trained in R2 still decoded frequency in R3 in the cue epoch and choice in the response epoch, respectively. This indicated that these representations were preserved after the rule switch (**fig. S2C)**. However, the two traces diverged in the response epoch and formed inverted, mirror-symmetric versions of each other. This pattern reflects the reversal of cue-action mappings from R2 to R3 and the gradual adaptation to the new rule, such that the projection from the sensory (cue frequency) subspace onto the action (choice) subspace was reoriented in the opposite direction (**fig. S2D**).

To quantify this reorientation, we measured cosine similarity between decoder hyperplanes in the second half of the response epoch (**Fig. 3C**). Alignment increased between cue location and choice in R1. In R2, alignment emerged between cue frequency and choice, which was then disrupted and reinstated in R3. Together, these results show that choice representations generalized across rules, whereas the alignment between the task-relevant cue dimension and the choice strengthened with learning and reorganized after rule switches.

### Stable activity and transient selectivity at the single-neuron level

Now we asked how single-neuronal responses changed across leaning. To distinguish the competing conceptual accounts (**Fig. 1**), we separately tracked each neuron’s temporal activity profile and its task-variable selectivity. To avoid selecting neurons by the activity we set out to characterize, we analyzed all 170 neurons recorded in all sessions rather than preselecting task-related units. To track the evolution of neuronal activity across learning, we sorted neurons by the latency of their activity in the R1 expert session (**Fig. 4A**). Latency was defined as the onset of five consecutive time windows (100 ms width), in which activity exceeded 80 % of the neuron’s peak averaged activity. No temporal sequence was evident in the novice session. A structured, trial-spanning pattern only formed gradually with learning (intermediate stage). This temporal structure remained stable across subsequent sessions and even persisted after rule switches. Next, we compared this activity pattern to the selectivity of task-relevant variables (cue location in R1, cue frequency in R2 and R3), preserving the neuron sorting by activity (**Fig. 4B**). In contrast to the stable activity profiles, the selectivity pattern lacked temporal structure and became increasingly fragmented and temporally unstable across sessions. Some neurons lost their selectivity, while others showed shifts in the timing of their peak selectivity.

**Fig. 4.**
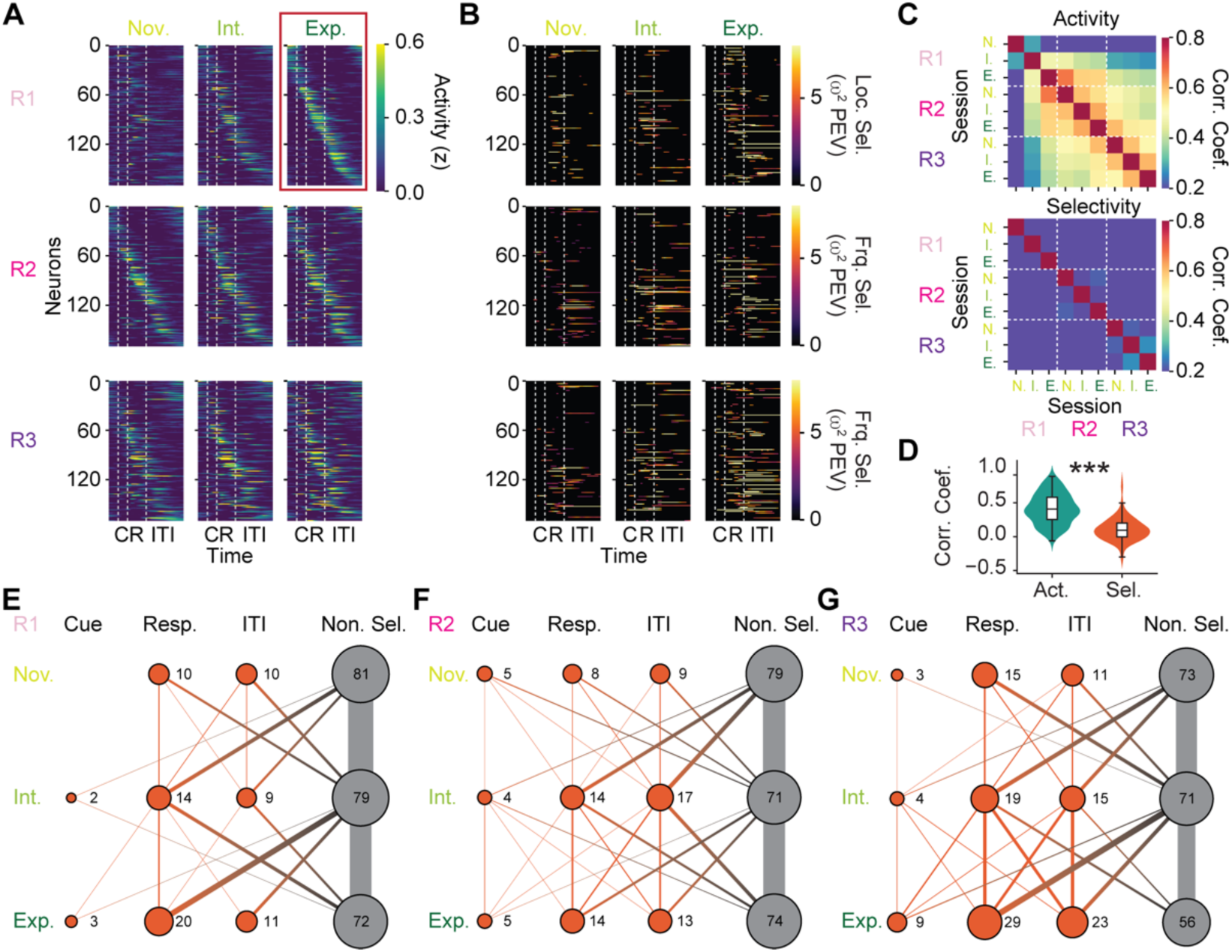
Activity-selectivity uncoupling across learning. (**A**) Mean calcium activity (z-scored) of all tracked neurons (*N* = 4 animals, *n* = 170 neurons), split by learning stages and rules. Neurons are sorted by activity latency in the R1 expert session (red box). (**B**) Selectivity (ω² PEV) for the task-relevant cue dimension (R1: location; R2, R3: frequency), preserving the same neuron order as in (A). (**C**) Pairwise Pearson correlation of single-neuron activity (top) and selectivity (bottom) between sessions, averaged across neurons. (**D**) Distributions of correlation coefficients in (C) across all neurons and session pairs (two-sample Kolmogorov-Smirnov test; ***, *p* < 0.001). Box, median and quartiles; whiskers, non-outlier range; points, outliers. (**E** to **G**) Percentage of neurons (total *n* = 402 (R1), 445 (R2), and 397 (R3) in *N* = 4 animals) selective for the task-relevant cue dimensions (R1: location; R2, R3: frequency), split by task epochs (cue, response and ITI) and learning stages. The links denote selectivity transitions between sessions. For each rule, all neurons were included that were recorded in all three sessions.

To quantify temporal stability, we computed Pearson correlations of activity and selectivity across sessions for each neuron. Activity correlations, averaged across neurons, were high overall (**Fig. 4C**, top). By contrast, the selectivity correlations exhibited near-zero off-diagonal values, reflecting poor consistency across sessions (**Fig. 4C**, bottom). Activity was significantly more stable than selectivity (**Fig. 4D**).

For each rule, we quantified the transition of selectivity at the single-neuron level by calculating the percentage of neurons selectively encoding the task-relevant cue dimension in different epochs (*n* = 402, *n* = 445, and *n* = 397 neurons recorded in all three sessions of R1, R2 and R3, respectively). The proportion of selective neurons increased across the entire training duration (Fig. 4E to **G**). Cue selectivity emerged only after the R1 novice session (**Fig. 4E**). Importantly, in all rules, the dominant transition pattern was between non-selective and selective neurons (31.0 % ± 1.3 of neurons, averaged across all six transitions). Shifts of selectivity between different epochs (7.4 % ± 1.8) and maintenance of selectivity in the same epoch across sessions (8.8 % ± 1.8) were much less frequent. In other words, distinct, non-overlapping, subsets of neurons were recruited at different learning stages. Together, these results support dynamic selectivity changes across learning at the single-neuron level, embedded within a stable prefrontal “activity frame” for task structure (activity-selectivity uncoupling; **Fig. 1B**, H_1_).

### Reconciling single-neuron transient selectivity with stable population representations

Next, we asked how the stable, low-dimensional population representations (**Fig. 3**) can be reconciled with the transient single-neuron selectivity and *ad hoc* recruitment across sessions (**Fig. 4**). Stability at the population level does not require neurons with stable tuning (*23*). If each representation occupies a distinct, stable subspace, a neuron’s contribution to the population code can be described using two properties (factors): first, its functional identity, i.e., which subspaces the neuron can engage in (that is, which representations a neuron can carry); and second, its expression, i.e., how strongly the neuron engages in the subspaces in a given session. In this framework, a neuron’s selectivity is then not an invariant intrinsic property, but rather the joint (multiplicative) result of a fixed functional identity and a session-varying expression. The observed loss and re-emergence of neuronal selectivity across sessions (**Fig. 4**) could therefore reflect changes in expression rather than a loss of identity. Within this framework, the same observed transient selectivity can arise in two ways. A neuron that engages in a single subspace (i.e., has a single identity) may do so more strongly in some sessions than others, appearing as selective when its expression is strong and non-selective when it is weak. Alternatively, a neuron that engages in several subspaces (i.e., has multiple identities) may shift how strongly it engages in each from session to session, re-weighting their combination and with it its observed selectivity.

Both accounts explain transient selectivity in terms of what individual neurons contribute to the population code, which is informative only if individual neurons are an identifiable part of that code, i.e., only if each representation can be traced to a sparse, strongly contributing set of neurons (*36*). In contrast, under a fully distributed, non-sparse code, every neuron instead contributes a very small share to every representation, meaning the population representation cannot easily be traced to individual neurons. Therefore, to make single-neuron contributions to the population code tractable, we decomposed population activity into task-related representations, each characterized by a specific temporal response motif (condition-specific activity; time factors), and implemented by a set of strongly coding neurons (**Fig. 5A**). Their contributions to each representation (loadings) are described by the product of neuron factors (functional identity) and session factors (expression) (**Fig. 5B**). Sparsity was imposed only on the neuron factors and not on the session factors, because neurons could in principle contribute to the same subspace persistently over time. Because this method operates on tensor-structured data, similar to tensor component analysis (*37*), we refer to it as Sparse Tensor Component Analysis (STCA).

**Fig. 5.**
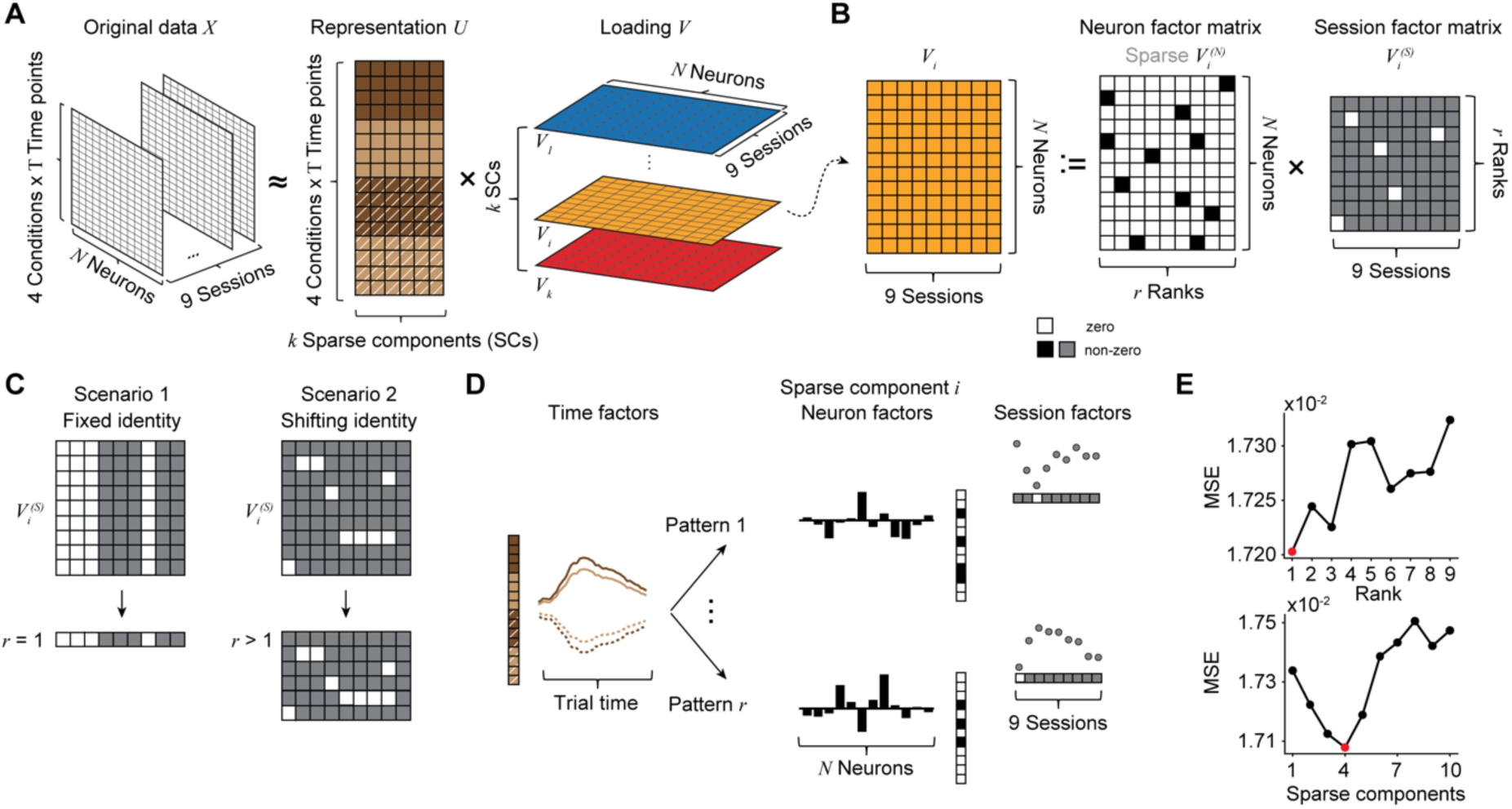
Sparse Tensor Component Analysis. (**A**) STCA factorizes the data tensor *X* into a representation matrix *U* and a loading tensor *V*; each tensor slice is the loading matrix of one sparse component (SC). (**B**) For SC *i* at rank *r*, the loading matrix factorizes into a neuron-factor matrix *V_i_*^(*N*)^ and a session-factor matrix *V_i_*^(*S*)^, with each rank defining a distinct neuron contribution pattern. Sparsity is imposed on the neuron-factor matrices. (**C**) Two scenarios for the session factor matrix regarding how a neuronal population contributes to an SC across sessions: fixed identity (rank 1) or shifting identity (rank > 1). The neuron factor matrix shares the same rank. Rows, contribution patterns; columns, sessions. (**D**) For each SC, a single temporal motif (represented by the time factors) can be carried by multiple neuronal subpopulations with distinct expression across sessions (represented by neuron factors and session factors, respectively). Neuron-factor vectors and session-factor vectors are separately constrained to be orthogonal across ranks. (**E**) Mean squared reconstruction error (MSE) between the demixed cue-dependent activity and its STCA reconstruction in the held-out test fold, plotted as a function of model hyperparameters (two-fold cross-validation); top, ranks per SC; bottom, number of SCs. Red dot, selected value.

STCA directly tests the above premise that a neuron’s functional identity is fixed. If fixed, each representation is carried by a single, stable subpopulation, yielding a rank-1 loading matrix (**Fig. 5C**, left). If instead a representation recruits different subpopulations at different times, the neuronal identity shifts across sessions and the loading matrix has higher rank (**Fig. 5C**, right). Transient selectivity would then reflect a shifting identity rather than the changing expression of a stable representation. The two possibilities can therefore be distinguished by the rank of the loading matrix for each sparse component (SC) (**Fig. 5D**).

We first isolated cue-dependent neuronal activity from time-dependent (i.e., cue-independent) signals and their interaction by demixing (*n* = 170 neurons recorded across nine sessions, see **Fig. 4**) (*38*). We then applied STCA to the cue-dependent activity and determined the number of SCs and their ranks using a grid search with two-fold cross-validation. The optimal model produced four SCs (reconstruction *R*^2^ = 0.159), each with a rank-1 loading matrix (**Fig. 5E**). These results confirmed that a neuron’s functional identity was indeed fixed, and that the observed transient selectivity instead reflected session-to-session changes in the extent to which it engages in a specific subspace.

### Neuronal engagement in multiple stable subspaces

SC time factors displayed condition-specific modulation, separated by either cue location (SC1) or cue frequency (SC2, SC3 and SC4) (**Fig. 6A**). These components were contributed by small subsets of neurons (high kurtosis of the neuron factor distribution, Sparsity Index (SI) > 1.0; Gaussian distribution: SI = 1) (**Fig. 6B**). Inspection of the session factors revealed that these sparse subpopulations were recruited in an *ad hoc*, rule-dependent manner: SC1 dominated in R1 (location rule), while SC2 dominated in R2 and R3 (frequency rules). SC3 and SC4 inverted their contributions from R2 to R3 (**Fig. 6C**). Neurons dominating each component (|*V*^(*N*)^| > 1.5 SD) showed stable activity, yet transient selectivity (**fig. S3**). Notably, behavioral performance was differentially reflected in each SC. SC1 and SC2 neurons showed higher activity in correct trials, while SC4 showed higher activity in error trials.

**Fig. 6.**
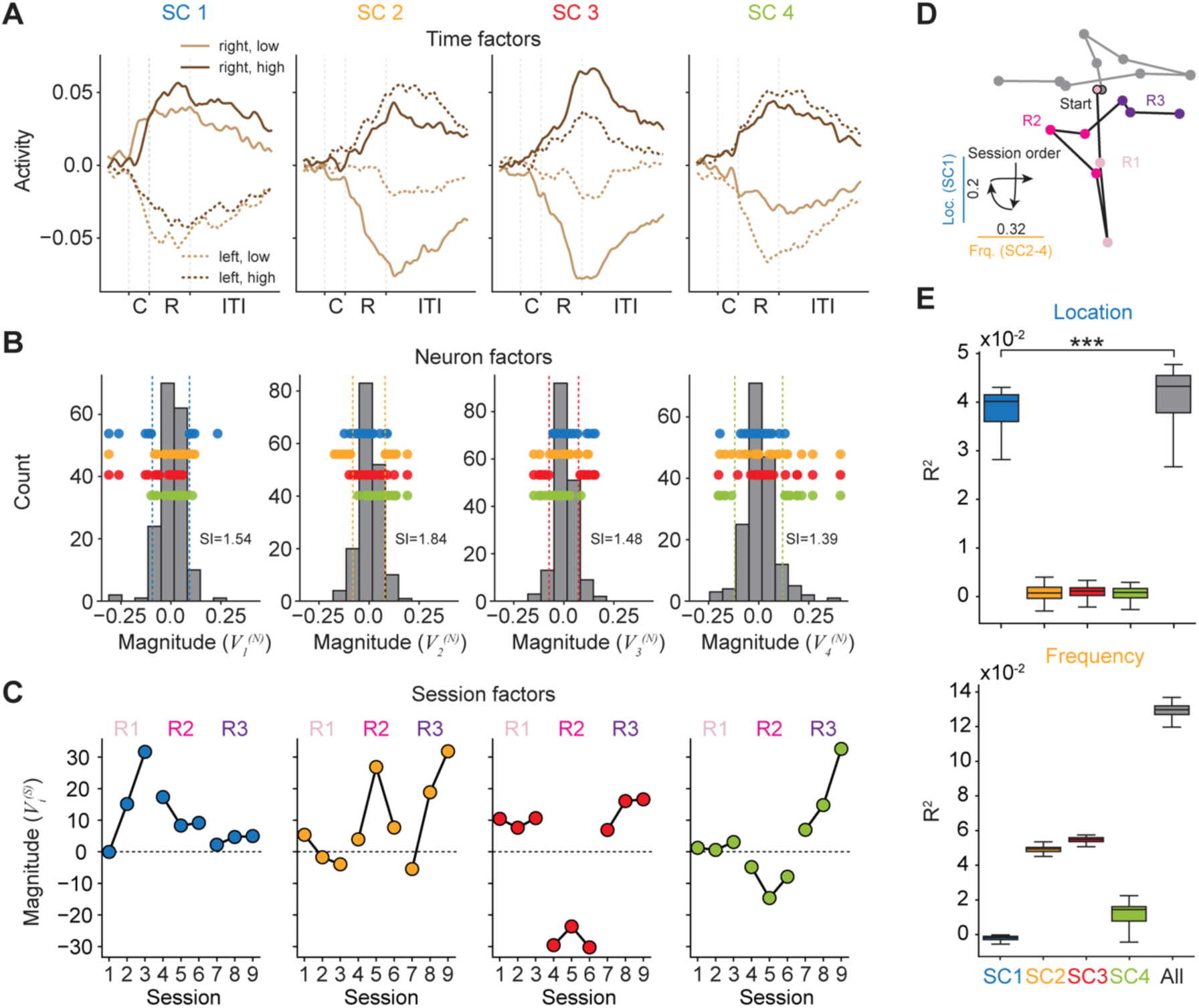
Subpopulations of multifunctional neurons. (**A**) Time factors for the four SCs and cue conditions. (**B**) Distributions of neuron factors ( *V_i_*^(*N*)^) for each SC. Colored dots mark dominant neurons of each SC (|*V_i_*^(*N*)|^> 1.5 SD, dashed lines). SI, sparsity index. (**C**) Session factors (*V_i_*^(*S*)^) for each SC. Signs for SC1, SC3, and SC4 are flipped for visualization. (**D**) Evolution of session-average neuronal loadings in a space spanned by SC1 and SC2-SC4. Two trajectories (color, gray) are shown for two neuron groups split by the sign of their SC4 neuron factors. Each dot marks one session. Arrows mark session order. (**E**) Prediction performance (*R*^2^) for cue location selectivity (top) and cue frequency selectivity (bottom) using single SCs or all SCs combined. Performance was evaluated by five-fold cross-validation across neurons, repeated 100 times. Box, median and quartiles; whiskers, non-outlier range; outliers not shown. ***, *p* < 0.001, Wilcoxon signed-rank test.

To investigate how rule-dependent recruitment evolved across learning, we projected loadings averaged across neurons in each session into a two-dimensional space spanned by SC1 (location contrast) and SC2–SC4 (frequency contrast) (**Fig. 6D**). The population state advanced along the location dimension during R1, shifted toward the frequency dimension after the rule switched to cue frequency (R2), and inverted along the frequency dimension once the cue–action mapping was reversed (R3).

Neuronal engagement in the SC subspaces was not mutually exclusive. Individual neurons contributed to multiple SCs (**Fig. 6B**), with an effective 2.2 ± 0.7 SCs per neuron (mean ± SD across 170 neurons). We refer to this property as neuronal multifunctionality. These neuron factors remained fixed across sessions, whereas each SC’s session factor determined the magnitude and sign of its expression in a given session. Changes in the relative session expression among SCs could therefore alter the net selectivity of a multifunctional neuron without changing its underlying functional identity. To test whether each neuron’s contributions to multiple SCs accounted for selectivity, we compared single-neuron ANOVA sum of squares (*SS*_Loc_ and *SS*_Frq_ ∝ *ω*^2^) with the log-transformed squared STCA-predicted condition contrasts for all individual SCs *SCs (△U_i_ = U^right^_i_ − U^left^_i_ or U^high^_i_ − U^low^_i_, i = 1, …, 4)* and their combination across sessions. STCA-predicted contrasts covaried significantly with measured selectivity for both cue dimensions (location, Spearman’s correlation *ρ* = 0.10, *p* < 10^-5^; frequency, *ρ* = 0.22, *p* < 10^-5^), indicating that STCA captured a significant portion of selectivity variability. To test whether multifunctionality is required to explain transient selectivity, we asked whether selectivity was better explained by a single SC or by their combination. For cue location, SC1 alone accounted for a substantial fraction of the variance, but the full model still performed significantly better (**Fig. 6E**, top). In contrast, under the frequency rules, no single SC explained selectivity well, whereas the combined model substantially improved prediction (**Fig. 6E**, bottom), indicating a stronger contribution of multiple SCs. Together, these results suggest that prefrontal neurons engage in multiple stable task-related subspaces and that the transient selectivity observed at the single-neuron level arises from re-weighting of their engagement in SCs across learning, rather than from changes of their functional identity.

## Discussion

We chronically recorded single-neuron activity in mPFC over several months as animals learned cue–action–outcome associations and adapted to successive rule changes. At the population level, task representations generalized across learning and tracked rule switches through the projection between them. At the single-neuron level, selectivity was strikingly transient. Neurons repeatedly gained, lost, or changed selectivity even after their temporal activity profiles had stabilized. STCA reconciled these observations by recovering a small set of population representations (SCs), each stably contributed by the same subset of neurons (its “substrate”), yet with neuronal engagement levels that varied from session to session. Thus, we identify a population-level, prefrontal cortical mechanism for cognitive flexibility, where volatility in single-neuron selectivity is not a destabilizing departure from a stable population code, but the expected signature of multifunctional neurons participating in task representations that are dynamically recombined as task demands evolve (**Fig. 7**).

**Fig. 7.**
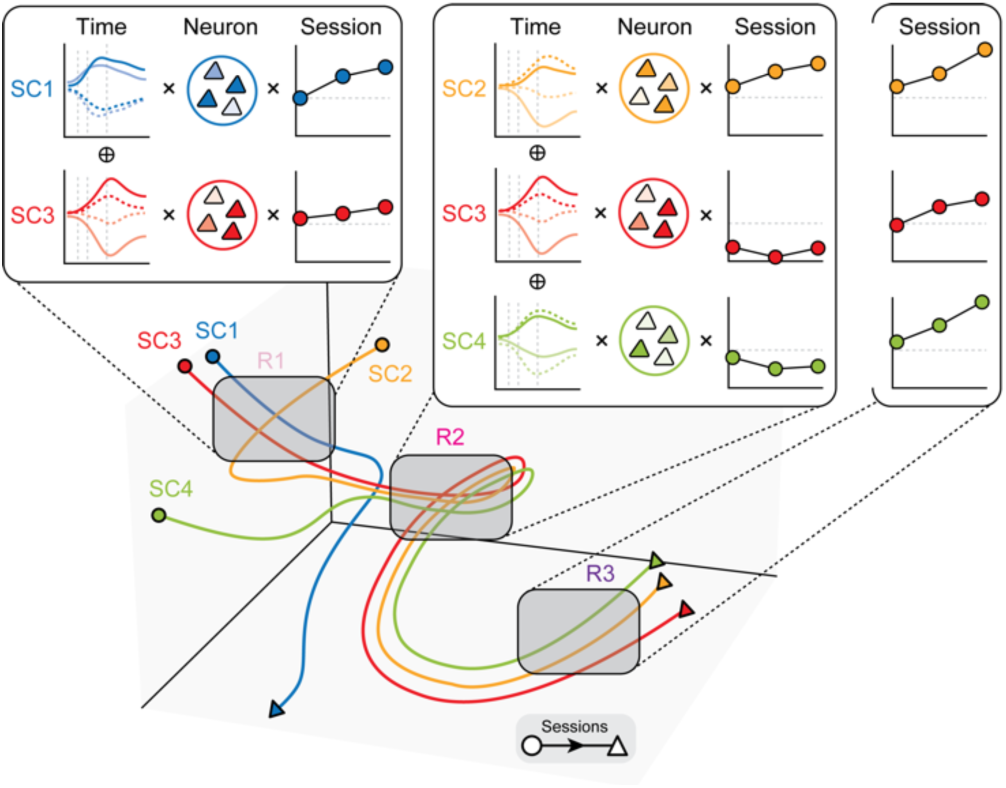
Rule-dependent subspace reconfiguration underlying transient selectivity. Schematic linking SCs to single-neuron selectivity. Population activity decomposes into four SCs, each a distinct representational subspace. Top, each SC contributes condition-specific activity (time factors) expressed differentially across sessions (session factors) and carried by specific neuronal subsets (neuron factors). Single-neuron selectivity arises from the weighted combination of these component-specific motifs. Multifunctional neurons engage in several SCs. Shifts in their SC engagement across learning results in transient selectivity. Bottom, state space trajectories show how different sets of SCs combine across learning and rules.

### Separating timing from content in prefrontal cognition

All three rules preserved the timing of task events, including cue presentation, behavioral responses and reward delivery after correct choices. Consistent with this common task structure, single-neuron activity tiled the trial at stable times across rules (**Fig. 4A**). This tiling is in line with the stable sequential dynamics previously reported in prefrontal cortex (*27–29*) and in thalamocortical circuits (*39, 40*), suggesting that temporally organized activity in PFC reflects an acquired stable scaffold of task structure (the “when”).

Within this scaffold, task-variable selectivity (the “what”) can be reconfigured, which is directly captured by STCA. The three rules imposed different cognitive computational demands. The R1-to-R2 switch required attention to a previously irrelevant dimension, in the sense of set shifting (*2, 5, 6, 41*). The R2-to-R3 switch did not change the relevant dimension but reversed the cue-action mapping, in the sense of reversal learning (*42, 43*). The organization of the SCs reflected this increase in computational complexity. Whereas the initial location rule was comparatively simple and dominated by a single SC (SC1), multiple SCs were required to capture the frequency contrast (**Fig. 6E**). The SC2-4 session factors captured signatures of these two learning demands. The interference from the previous location rule showed in the inverted U-shaped traces in R2 (peaking in the intermediate learning stage). After reversal and without extra-dimensional interference, the traces increased monotonically again in R3 (**Fig. 6C**).

### Relation of STCA to other linear dimensionality reduction methods

Existing dimensionality reduction methods do not capture the full structure recovered by STCA. They either decompose population activity without isolating a motif from the sparse set of neurons that express it, e.g., demixed PCA or tensor component analysis (*37, 38, 44*), or depend on predefined task labels and overlook latent motifs, e.g., targeted dimensionality reduction (*45, 46*). STCA addresses these limitations by imposing sparsity directly on neuron factors and extracting components without task labels, thereby isolating each motif from a small set of strongly contributing neurons while leaving their functional identity to be inferred from its within-trial dynamics. STCA therefore provides an unsupervised method for studying the computational neuronal organization of cognitive flexibility, in which an inventory of stable, low-dimensional motifs is recombined to implement distinct computations as task demands change (*46–48*).

### Multifunctional neurons and the substrate of flexibility

Mixed selectivity, where neurons respond to nonlinear combinations of task variables, is thought to support flexible computation by expanding the dimensionality of the population code beyond the number of task variables, so that arbitrary task variable contrasts can be linearly read out from the same neurons (*49–51*). Our results extend the mixed-selectivity framework in two ways. First, the relevant axes of mixing we identified represent integrative functional identities, which are extracted directly from population activity and captured as SCs. Traditional accounts of mixed selectivity, by contrast, are based on supervised, observer-defined isolated task variables. Second, this mixture is not static. A neuron’s selectivity in a given session can be approximated as a linear combination of its stable contributions to multiple SCs. Session factors determine when an SC is expressed in the population code, whereas neuron factors define the stable substrate that supports it. Task variable selectivity can therefore change substantially across sessions even when a neuron’s functional identity, its engagements in SCs, remains fixed.

### Dynamic recruitment as an adaptive strategy

Multifunctionality accounts for only part of the volatility in single-neuron selectivity (**Fig. 6E**), and additional sources likely contribute. From the learning animal’s perspective, the task is not a fixed external structure but an internal model it continually infers and re-parameterizes (*7*). As learning proceeds, policy changes (*52, 53*), credit assignment shifts (*54, 55*), and temporal compression can reshape the computation (*56*), rotating the relevant subspaces and varying the mapping between neurons and those subspaces (*45, 46*). Such dynamics preserve low-dimensional motifs and decision geometry while changing which neurons most strongly instantiate them across sessions. Because this task-belief updating across learning is continuous rather than discretely segmented into separable stages, the mapping between neurons and subspaces cannot settle into a final form. Under activity-selectivity uncoupling (H_1_), plastic neuron-to-subspace mappings can reassign neuronal selectivity and update internal variables as animals adapt. By contrast, under activity-selectivity coupling (H_0_), selectivity is bound to activity, so the same update would collapse the activity scaffold and require a rebuild, discarding what had been learned rather than keeping it for reuse. The transient selectivity coexisted with a stable, decodable population representation that tracked rule switches (**Fig. 3**), suggesting it is a crucial feature of the prefrontal code rather than a limitation of it.

### The neuronal circuits of cognitive flexibility

Our analyses identify a computational organization but do not reveal its underlying circuit architecture. One possibility is that the stable components identified by STCA are implemented through the flexible recruitment of projection-defined neuronal populations. Supporting this possibility, striatum-projecting orbitofrontal neurons can selectively and persistently encode an individual decision variable (*57*), while dorsomedial-striatum-projecting mPFC neurons form temporally structured populations and contribute specifically to working-memory maintenance (*58*). Combining longitudinal recordings during learning with projection-specific labeling and perturbation will be necessary to determine whether dynamic recruitment reflects the selective engagement of anatomically defined output pathways.

## Acknowledgments

We thank X.-X. Lin, J. Gjorgjieva, M. Stemmler, and members of the Jacob Lab for insightful discussions. Grammarly, ChatGPT, and Claude were used in language editing and drafting supporting. No AI tools were used in data interpretation.

## Author contributions

Conceptualization: YYH, LSM, SNJ

Methodology: YYH, LSM, TWB, SNJ

Investigation: LSM

Data curation: YYH, LSM

Formal analysis: YYH

Software: YYH

Visualization: YYH

Funding acquisition: LB, SNJ

Project administration: SNJ

Supervision: LB, SNJ

Writing – original draft: YYH, SNJ

Writing – review & editing: YYH, LSM, LB, SNJ

## Data, code, and materials availability

All data, code, and materials are available from the corresponding author.

## Materials and Methods

### Animals

All procedures were approved by the local regulatory authority (Regierung Oberbayern). Male wild-type mice (C57BL/6J, Charles River), aged 8–10 weeks at the start of the experiments, were housed individually under a reversed 12-hour light/dark cycle (lights off during the day). Temperature and humidity were maintained at 24°C and 50%, respectively. Mice had unrestricted access to food and water, except during behavioral sessions. Animal health was monitored and scored daily.

### Behavior task

Mice were pre-trained to respond to licking spouts by receiving three to four sessions where only one spout was presented per trial in random order, with a water reward delivered after a correct lick. Once mice consumed at least 800 µl of water in a session, they advanced to the main task. In the auditory decision-making task, all 12 implanted mice learned to lick one of two spouts for a water reward (5 µl) based on an auditory instruction cue: either a low-frequency (4–8 kHz) or high-frequency (16–32 kHz) bandpass-filtered white noise at 75–80 dB. Each trial began with a luminance increase on a monitor, followed by the auditory cue (1,000 ms) and the presentation of both spouts. The first tongue contact during the 2,000-ms response window was registered as the response. Correct licks triggered reward delivery and retraction of the non-target spout. Incorrect licks led to retraction of both spouts and a 4,000-ms timeout. In miss trials (no lick), both spouts were retracted after the response window.

Animals were trained on three implicit task rules based on cue location and frequency. First, in the location rule, mice learned to respond to the left or right spout based on which speaker played the cue. Once animals had acquired the first rule, the rule was switched to the next. Then, in the frequency rule, responses were based on the cue’s frequency. Finally, the cue-action mapping of the high and low frequencies was switched in a reversed frequency rule. In a fraction of trials, only the target spout was presented, forcing the animal to make a correct choice. These no-choice (forced) trials were interleaved within every session and across all task rules. Because learning was modestly slowed after prelimbic cortex implantation, they aided rule acquisition and kept responding balanced across the two spouts, reducing side biases. Forced trials were excluded from analysis.

### In vivo chronic calcium imaging

Chronic in vivo calcium imaging was performed using head-mounted miniature microscopes (UCLA Miniscopes) to monitor neuronal activity. GRIN lenses (0.5 mm diameter, ∼6.1 mm length; Inscopix) were pre-coated with a mixture of silk fibroin and AAV (AAV1.CamKII.GCaMP6f.WPRE.SV40, Addgene) (*59*). After a craniotomy above the prelimbic mPFC, the coated lens was slowly lowered to the target depth, delivering the virus during implantation. The implant was secured to the skull with dental adhesive and cement. After recovery and at least four weeks of viral expression, a miniscope baseplate (Miniscope Version 3, UCLA) was attached in a subsequent, non-invasive procedure to allow repeated imaging. Imaging sessions were conducted during three stages (novice, intermediate, and expert performance) in each rule, and calcium signals were acquired at a frame rate of 30 Hz. Mice were habituated to head-mounting before recording. The novice sessions were defined as the first session of each rule, while the expert sessions were recorded after mice reached a 70 % correct rate. The intermediate sessions were recorded when the performance was between 60 % and 70 %.

### Histological processing

Following behavioral and calcium imaging experiments, animals were perfused with 4 % paraformaldehyde. Brains were post-fixed for 24 h with miniscopes in place, then sectioned coronally at 120 µm using a vibratome. Sections were covered with mounting medium (VectaShield) and imaged using a confocal microscope (Leica SP8) with tenfold magnification. Confocal images were aligned with the mouse brain atlas (*60*) for verification of implantation locations.

### Extraction of calcium signals

Neuronal calcium signals were extracted using the open-source Python package CaImAn (*33*). As an initial step for motion correction, the algorithm applies spatial high-pass filtering using a Gaussian kernel with a size of 7 pixels to suppress background and enhance structural features. Motion correction is then performed using the NoRMCorre algorithm in CaImAn to infer the rigid shifts of each frame, and the estimated motion is applied to the original data to preserve signal integrity. Source extraction was conducted using the CNMF-E algorithm with parameters (minimum correlation of 0.8 and peak-to-noise (PNR) ratio of 10) empirically selected. The correlation threshold was chosen around the local minimum in the bimodal correlation distribution, and the PNR threshold was determined around the 70th percentile. Regions of interest were evaluated based on their spatial correlation to active frames (rval) and signal-to-noise ratio (SNR), with only components surpassing quality thresholds retained. Components with rval > 0.85 and SNR > 3 were kept for further analyses.

### Longitudinal registration of active neurons

We utilized an in-built CaImAn function to chronologically register ROIs of the current session against the union of units of all the past sessions aligned to the current field of view (FOV). It first calculates a pairwise distance matrix measuring the overlap ratio between any two binarized ROIs that are close enough in two aligned FOVs. Then it solves the linear assignment problem using Hungarian algorithm for the best match. This method allows components from both sessions that are not matched with any other component to prevent false assignments and enables the registration of sessions with different numbers of neurons. The threshold for the overlap ratio (thresh_cost) was set to 0.7, and the threshold of centroid distance for calculating the overlap ratio (max_dist) was set to 20 pixels. By examining the quality metrics, we found that units detected in more than one session have significantly better quality than those detected only in one session (SNR, rval, **fig. S1**). Therefore, only units detected in more than one session were retained for further investigation.

### Single-neuron selectivity (Fig. 2I, 3B)

To assess the selectivity of individual neurons over time, we performed a two-way ANOVA on cue location and frequency in a sliding window across the trial. For each neuron, calcium activity traces were segmented into fixed-length temporal windows (400 ms) with a defined step size (200 ms). Within each window, the mean activity was computed. We used two-way ANOVA to assess the effect of task variables and their interaction on neuronal activity. The significance threshold was set at family-wise error rate < 0.05 (Bonferroni-corrected for multiple comparisons). A neuron was considered selective if any main effect was significant in any session. For population-average selectivity and correlation across sessions, we used effective selectivity quantified by subtracting the chance-level selectivity estimated from trial-shuffled data using permutation testing. Trial labels were randomly shuffled 100 times, and two-way ANOVA was recomputed for each permutation and time window. This generated a null distribution of selectivity values for each factor. The 99^th^ percentile of the corresponding null distribution was taken as the chance-level selectivity at each time window.

### Cross-temporal decoding (Fig. 3A, fig. S2A)

We performed cross-temporal decoding using a linear classifier (support vector machine, SVM, sklearn.svm.SVC) per animal. Neural activity was binned into sliding windows (200 ms width, 200 ms step). At each time point, a classifier was trained to discriminate conditions (cue location, frequency, or behavioral choice) using activity patterns across neurons. The trained classifier was then tested on all other time points to generate a cross-temporal decoding matrix, where each element reflects the decoding accuracy when trained at time *t*_train_ and tested at time *t*_test_. To maintain statistical power in novice sessions, we included both correct and error trials in the analysis; however, this inclusion did not qualitatively affect the results (fig. S2A).

An equal number of trials per condition was subsampled to ensure balanced class representation in each training set. Classification performance was evaluated using five-fold cross-validation, repeated across folds to obtain the average accuracy. This procedure was repeated 100 times, and the final decoding matrix for each animal was computed by averaging across all repetitions. Statistical significance was assessed via a permutation test. Trial labels were randomly shuffled (100 permutations), and decoding was recomputed to generate a null distribution for each matrix element. The 25^th^ and 75^th^ percentiles were used to mark significant deviations.

### Cross-session decoding (fig. S2B, C)

The cross-session decoding takes a similar procedure to the cross-temporal decoding, except that classifiers (for cue location, frequency, or behavioral choice) were tested in the same time window (200 ms width, 100 ms step) in other sessions using all balanced trials. The final decoding accuracy for each animal was averaged across 100 repetitions.

We calculated the cosine similarity between their normal vectors to quantify the alignment between classifier hyperplanes within the same session. Each normal vector was derived by averaging the weight vectors of a linear classifier trained over 100 repetitions. Because the hyperplanes are defined in the same neuronal activity space, their orientations are directly comparable. For comparisons across sessions, we first identified overlapping neurons between any two sessions, concatenated their activity, and applied principal component analysis (PCA). The hyperplane normal vectors were then computed using the first three principal components to reduce noise in population responses. Trial numbers were balanced across conditions in all analyses.

### Sparse tensor component analysis (Fig. 6A-C)

We bridge single-neuron activity with population representation by introducing Sparse Tensor Component Analysis (STCA), which overcomes a key limitation of the previous Sparse Component Analysis (SCA) framework (*36*): the inability to track neuronal contributions across long-term recordings. Each component (SC) consists of a temporal motif, a set of neuron factors, and session factors reflecting its population-level expression. We did not impose sparsity on the session factors, as neurons could, in principle, contribute to task representations persistently across sessions.

The resulting STCA incorporates session-specific factors into the dimensionality reduction process, capturing how neuronal contributions to representations evolve across different learning stages. Following the formulation of SCA, we determine the *k* sparse components in the neuronal activity *X* of *N* neurons with trial time length *T* under *C* conditions across *S* sessions based on the linear generative model

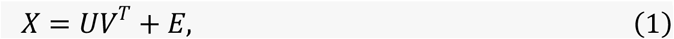

where *X* ∈ ℝ*^CT^*^×*NS*^is approximated by *k* firing activity vectors *U* ∈ ℝ*^CT^*^×*k*^ and their corresponding loading *V* ∈ ℝ*^NS^*^×*k*^, with columns denoting the components. Here, *E* denotes the residual matrix. The loading on each sparse component *V_i_* is defined as

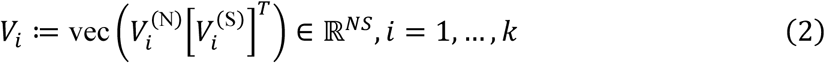

where:

- *V*^(N)^ ∈ ℝ*^N^*^×*r*^ is the neuron factor matrix with rank *r*,
- *V*^(S)^ ∈ ℝ*^S^*^×*r*^ is the session factor matrix with rank *r*,
- vec(⋅): ℝ*^d^*^1×*d*2^ → ℝ*^d^*^1^*^d^*^2^ flattens a matrix into a vector.

Interpreting *U_i_* as a representation and *V_i_* as its neuronal implementation, each entry of *V_i_* reflects the contribution of a given unit in a specific session. If *V_i_* ∈ ℝ*^N^*^×*S*^has a rank of 1, it can be expressed as the outer product of a column vector *v_i_*^(N)^ and a row vector *v_i_*^(S)^, indicating that one single neuronal subpopulation contributes dynamically across sessions. In contrast, a rank greater than 1 implies the involvement of multiple subpopulations, each with distinct contribution patterns over sessions. Thus, the rank of the neuron and session factor matrices determines both the number of contributing subpopulations to the representation and the variability of their contributions across learning stages.

Compared with SCA, STCA seeks the optimal components and their neuronal implementations by enforcing the sparse penalty on the neuron factor tensor *V*^(N)^ ∈ ℝ*^k^*^×*N*×*r*^

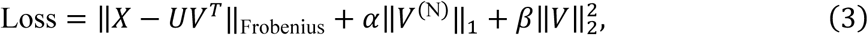

instead of applying the penalty on the loading *V* itself, as we only assume a neuron sparsity. Here, ‖ ⋅ ‖*_p_* denotes the *p*-norm. The parameter *α* controls the extent of sparsity constraint on neuron factor via *L*_1_ regularization, and *β*, set to 0.1, smoothes the loss landscape and stabilizes the results across random initialization. The parameters, *α*, *k*, and *r*, were determined by grid search using two-fold cross-validation. The sparsity of each component is quantified by Sparsity Index (SI), which is proportional to the kurtosis of the neuron factor distribution and equal to 1 for a Gaussian distribution (*36*).

### Demixing for STCA

We demixed the activity of *n* = 170 neurons, which we could follow across *S* = 9 imaging sessions, from four mice for STCA. We first consider an individual neuron in a single session. The activity matrix *x* can be decomposed into marginalizations over two parameters, time (*t*) and cue (*c*) (*36, 38*)

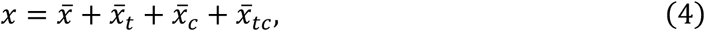

The cue-dependent part is then the cue marginalization grouped by its interaction with time, because the cue is modulated by time, i.e.,

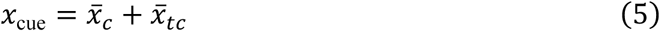

The resulting activity matrix for all neurons is then *X* ∈ ℝ*^C^*^×*N*×*T*^.

The trial-wise residuals (trial activity minus the condition mean) were used to calculate the across-trial standard deviation for each neuron and time point. Condition means were then z-scored by dividing by this standard deviation. To prevent numerical instabilities, undefined values resulting from zero variance were replaced with zeros. Conditional means from 9 sessions were stacked to form a matrix *X* ∈ ℝ*^S^*^×*C*×*N*×*T*^, which was then reshaped to ℝ*^CT^*^×*NS*^for input to STCA.

### Dynamics in implementation space (Fig. 6D)

We grouped neurons into two categories based on the sign of their neuron factor in SC4. We chose SC4 because its session factor reversed between R2 and R3 and its least sparsity index reflected broad neuronal involvement, giving a reliable basis for defining coding polarity across the population. For each group, we calculated contributions to each SC in each session by averaging their neuron factors and multiplying by the corresponding session factor. We then obtained the scalar sum of contributions from the frequency-modulated SCs (SC2–SC4) and compared these with the contributions along the location-modulated SC (SC1).

### Quantification of neuronal multifunctionality

To quantify the degree to which individual neurons participated in more than one sparse component (SC) without imposing a loading threshold, we analyzed the neuron-mode factors of the four-component decomposition, arranged as a matrix *A* of 170 neurons × 4 SCs. Because the decomposition identifies each component only up to a reciprocal rescaling of its neuron and session factors—the fitted temporal factors were constrained to unit norm whereas the neuron factors were not (column norms 0.64–1.05)—the absolute magnitude of a neuron factor is not comparable across components. We therefore L2-normalized each component’s neuron-factor vector across neurons (dividing each column by its Euclidean norm, without mean-centering) so that the four SCs contributed on a common scale; results were confirmed to be insensitive to this step (Spearman ρ = 0.86 between normalized and unnormalized values). For each neuron *n* we computed the proportional squared contribution of SC *i*,

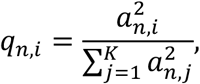

and summarized participation as the effective number of SCs,

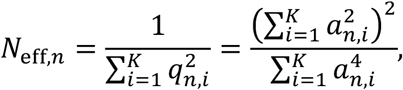

the participation ratio (inverse Simpson index) of the squared factor magnitudes. Squaring the factors ensured that positive and negative loadings did not cancel, and every factor magnitude was retained (no threshold was applied). *N*_eff,*n*_ ranges from 1, when a neuron’s factor magnitude lies entirely in a single SC, to 4, when it is distributed equally across all four SCs, with intermediate values indicating graded multifunctionality; it is invariant to sign flips and to rescaling of a neuron’s factor vector and was treated as undefined for any all-zero neuron (none occurred). Statistics were computed descriptively across the pooled population of all 170 neurons, reported as mean ± SD and median with interquartile range. A 95 % confidence interval for the mean was obtained by a neuron-level bootstrap (10,000 replicates resampling neurons with replacement; fixed seed for reproducibility). Because neurons could not be attributed to individual animals in this dataset, the reported dispersion describes heterogeneity across the recorded neurons and the confidence interval is conditional on this pooled sample rather than an estimate of between-animal variability.

Correlation analysis between STCA contrast and ANOVA selectivity (Fig. 6E)

To quantify the association between STCA-predicted contrast and ANOVA selectivity, we derived factor-specific contrast predictors from the STCA-reconstructed condition means. For each sparse component *i*, reconstructed temporal motifs were first averaged across conditions sharing the same task variable. For cue location, the component-level contrast was defined as

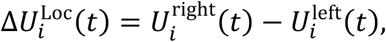

where *U^right^_i_(t)* and *U^left^_i_(t)* denote the temporal motifs associated with right- and left-cue conditions, respectively. Analogously, the frequency contrast was defined as

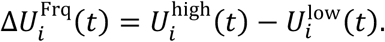

For neuron *n* in session *s*, the predicted contrast was then computed as the weighted sum of component-wise contrasts across all sparse components,

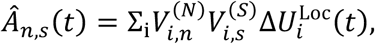

and

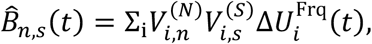

where *V^(N)^_i,n_* and *V^(S)^_i,s_* denote the neuron and session factors of component i, respectively. Squared predicted contrasts *Â^2^_n,s_(t), B^^2^_n,s_(t)* were compared with ANOVA sums of squares (*SS_A_*, *SS_B_*), since sums of squares scale with squared condition contrasts in balanced designs. We used sums of squares rather than ω² because they scale directly with the squared condition contrast that STCA predicts, whereas ω² additionally divides that contrast by each neuron’s total variance, which STCA does not predict.

For each retained unit, session, and sliding-window time bin, we paired the observed cue sum of squares from the 2-way ANOVA with the model-based contrast predictors. Location and frequency were analyzed separately: log(1 + SS_Loc_) with log(1 + *A*^^2^), and log(1 + SS_Frq_) with log(1 + *B*^2). Observed SS and predictors were stored as three-dimensional arrays (neurons × sessions × time bins) and flattened into paired one-dimensional vectors over all neuron–session–time entries. Spearman’s rank correlation coefficient *ρ* and the associated two-sided *p*-value were computed with *scipy.stats.spearmanr* on these paired vectors.

### Compositional Contrast Prediction Test (Fig. 6E)

We evaluate whether ANOVA selectivity is better predicted by single contrast components or by their superposition by performing five-fold cross-validation with neurons as the split unit: for each fold, we train linear models on one subset of neurons and test on held-out neurons after flattening session and time dimensions, fitting separate univariate regressions for each squared single-component predictor and one regression for the squared combined predictor, then averaging test across folds to obtain per-component and combined performance; this procedure is repeated 100 times with different random seeds to estimate stability.

---

**fig. S1.**
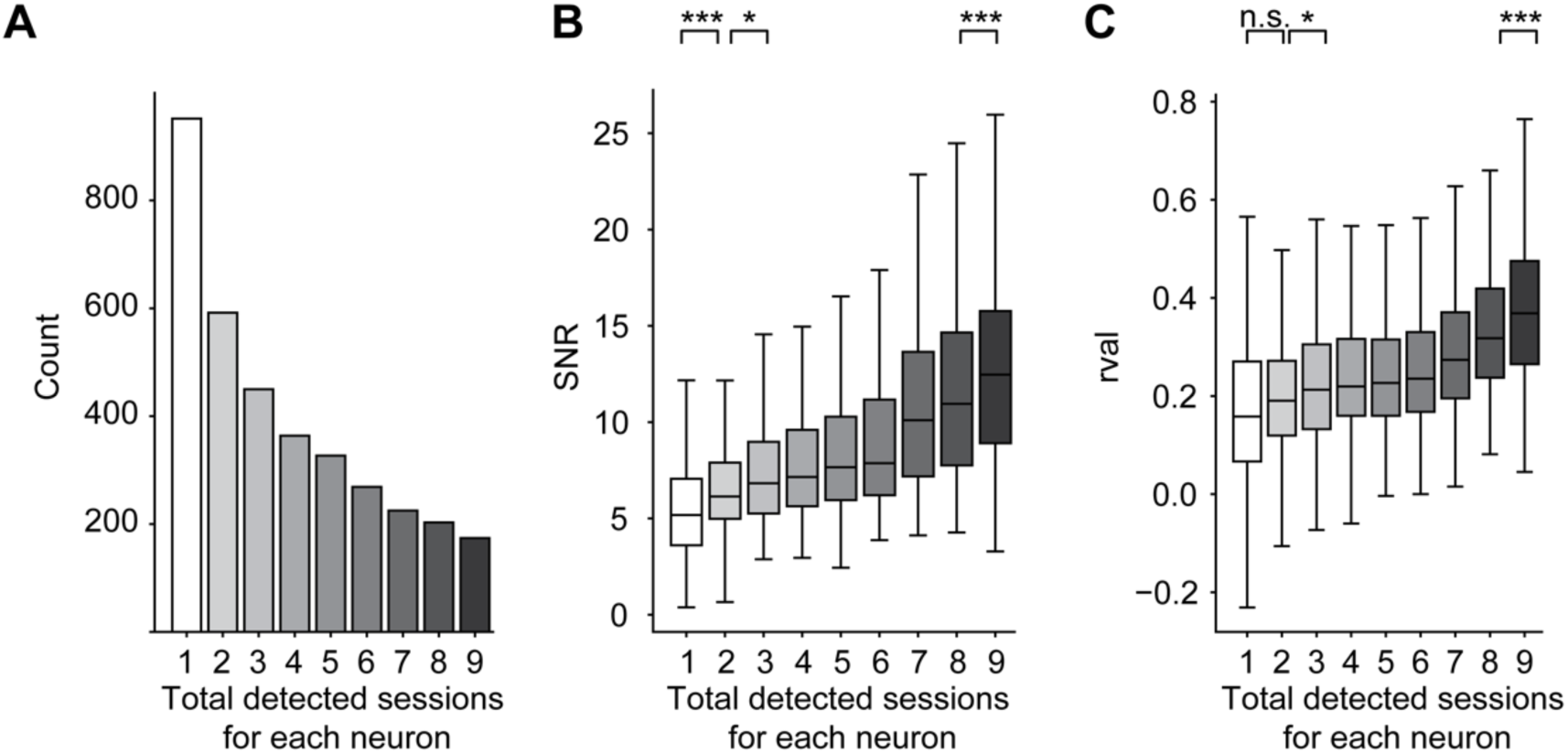
Calcium imaging quality metrics. (**A**) Neuron count split by the number of sessions in which each was detected (*N* = 12 animals). (**B**) Same layout as in (A) for signal-to-noise ratio (SNR; the likelihood of the measured trace under an estimated noise distribution). Significance from a mixed-effects model (SNR∼session + (1|mouse): *p*_12_ = 1.2 × 10^-4^, *p*_23_ = 1.3 × 10^-2^, *p*_89_ = 8.5 × 10^-5^). (**C**) Same layout as in (A) for spatial footprint consistency (rval; the correlation of each spatial footprint with its own average active frame). Significance from a mixed-effects model ( rval∼session + (1|mouse): *p*_12_ = 8.8 × 10^-2^, *p*_23_ = 2.7 × 10^-2^, *p*_89_ = 4.2 × 10^-4^). Box, median and quartiles; whiskers, non-outlier range; outliers not shown. *, *p* < 0.05, **, *p* < 0.01, ***, *p* < 0.001; n.s., not significant.

**fig. S2.**
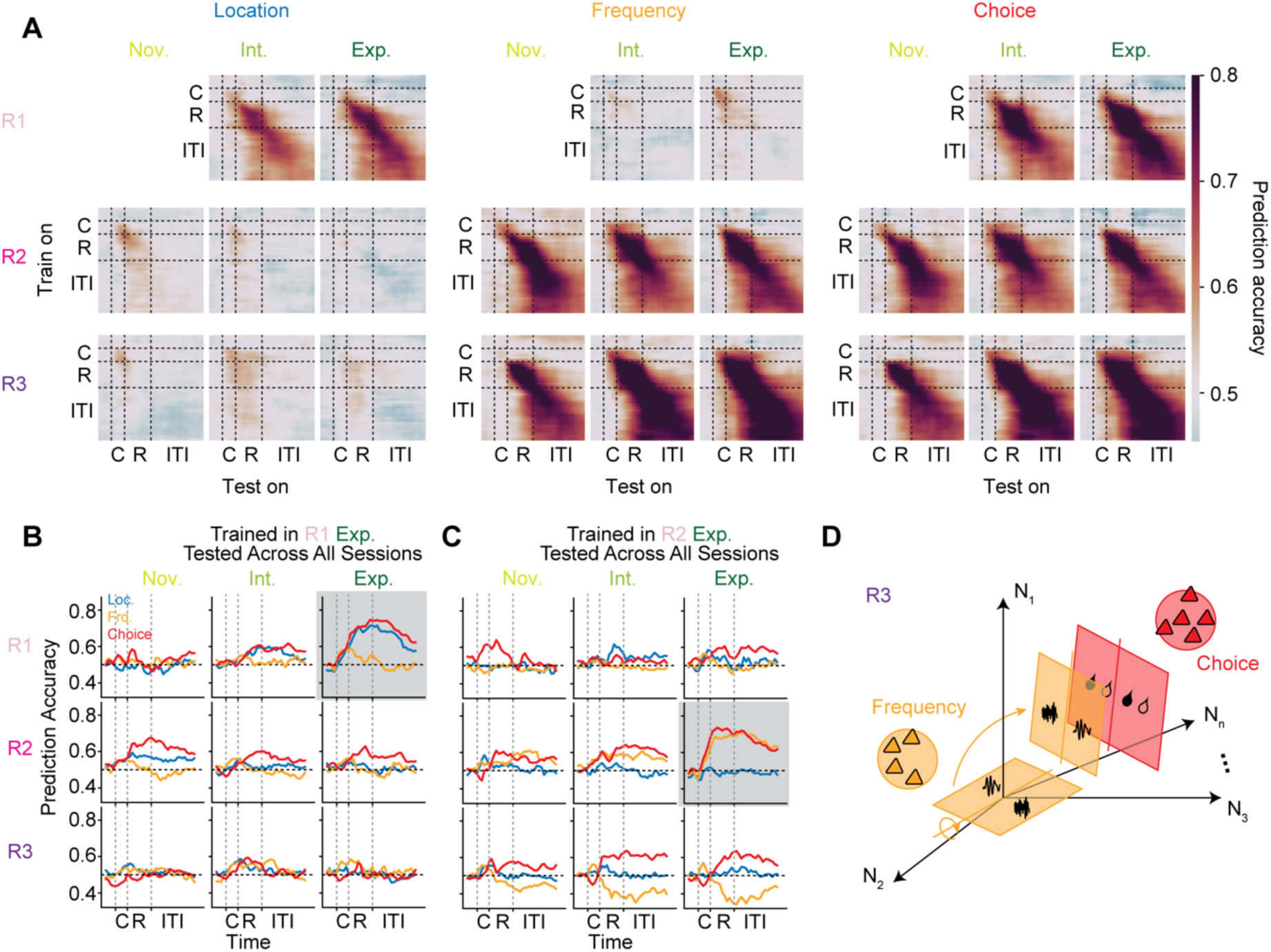
Reorganization of population representations. (**A**) Cross-temporal decoding as in Fig. 3A but using correct trials only (*N* = 4 animals, *n* = 149 neurons); R1 novice is omitted (too few correct trials per condition). (**B**) Cross-session decoding. Binary SVM classifiers trained in the R1 expert session (shaded) and tested at the same trial time across all sessions, decoding cue location, cue frequency, and behavioral choice (*N* = 4 animals, *n* = 149 neurons; correct and error trials). (**C**) Same layout as in (B), but trained in the R2 expert session (shaded). (**D**) Schematic of subspace transformation after the rule switch to R3. The frequency subspace rotates to its inversion after cue presentation, reflecting a reversal of the cue-action mapping.

**fig. S3.**
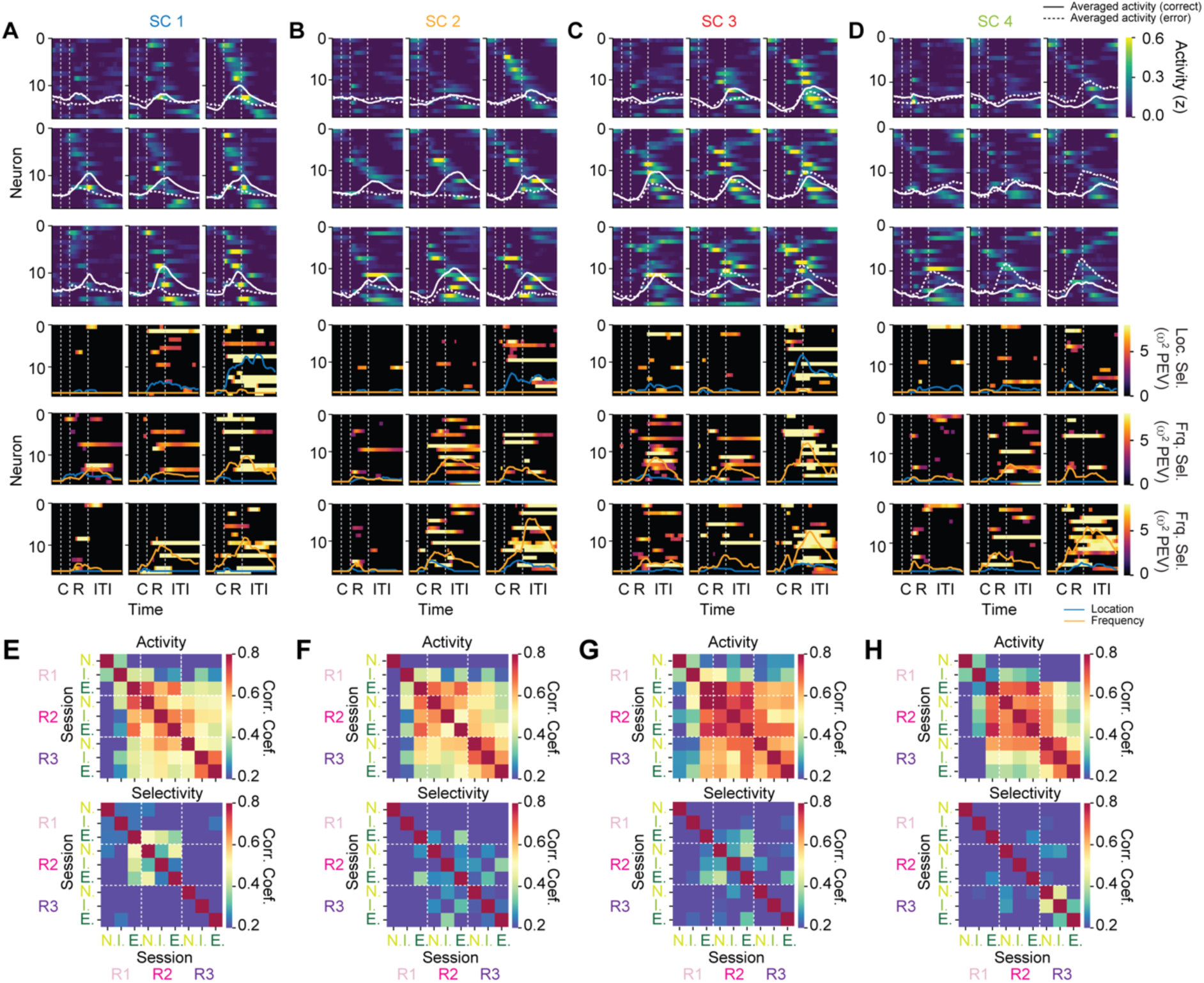
Activity and selectivity of SC dominating neurons. (**A**) Top, mean calcium activity (z-scored) of the dominant neurons for SC1, split by learning stages and rules. Neurons are sorted by activity latency in the R1 expert session (red box). Trial-averaged activity is overlaid (solid, correct trials; dashed, error trials). Bottom, selectivity (ω² PEV) for the task-relevant cue dimension (R1: location; R2, R3: frequency), preserving the same neuron order as above. (**B** to **D**) Same layout as in (A) for SC2, SC3 and SC4. (**E**) Pairwise Pearson correlation of single-neuron activity (top) and selectivity (bottom) between sessions, averaged across the dominant neurons of SC1, as in Fig. 4C. (**F** to **H**). Same layout as in (E) for SC2, SC3 and SC4. Dominant neurons as defined in Fig. 6B.

## References

1. M. E. Ragozzino, S. Detrick, R. P. Kesner, Involvement of the Prelimbic-Infralimbic Areas of the Rodent Prefrontal Cortex in Behavioral Flexibility for Place and Response Learning. J Neurosci 19, 4585–4594 (1999).

2. D. Durstewitz, N. M. Vittoz, S. B. Floresco, J. K. Seamans, Abrupt transitions between prefrontal neural ensemble states accompany behavioral transitions during rule learning. Neuron 66, 438–448 (2010).

3. M. P. Karlsson, D. G. R. Tervo, A. Y. Karpova, Network Resets in Medial Prefrontal Cortex Mark the Onset of Behavioral Uncertainty. Science 338, 135–139 (2012).

4. D. R. Euston, A. J. Gruber, B. L. McNaughton, The role of medial prefrontal cortex in memory and decision making. Neuron 76, 1057–1070 (2012).

5. T. Spellman, M. Svei, J. Kaminsky, G. Manzano-Nieves, C. Liston, Prefrontal deep projection neurons enable cognitive flexibility via persistent feedback monitoring. Cell 184, 2750–2766 e2717 (2021).

6. A. Del Arco, J. Park, J. Wood, Y. Kim, B. Moghaddam, Adaptive Encoding of Outcome Prediction by Prefrontal Cortex Ensembles Supports Behavioral Flexibility. J Neurosci 37, 8363–8373 (2017).

7. R. Bartolo, B. B. Averbeck, Prefrontal Cortex Predicts State Switches during Reversal Learning. Neuron 106, 1044–1054 e1044 (2020).

8. N. J. Powell, A. D. Redish, Representational changes of latent strategies in rat medial prefrontal cortex precede changes in behaviour. Nat Commun 7, 12830 (2016).

9. S. Reinert, M. Hubener, T. Bonhoeffer, P. M. Goltstein, Mouse prefrontal cortex represents learned rules for categorization. Nature 593, 411–417 (2021).

10. E. M. Meyers, X. L. Qi, C. Constantinidis, Incorporation of new information into prefrontal cortical activity after learning working memory tasks. Proc Natl Acad Sci U S A 109, 4651–4656 (2012).

11. M. Sakagami, H. Niki, Spatial selectivity of go/no-go neurons in monkey prefrontal cortex. Experimental Brain Research 100, 165–169 (1994).

12. J. D. Wallis, K. C. Anderson, E. K. Miller, Single neurons in prefrontal cortex encode abstract rules. Nature 411, 953–956 (2001).

13. L. Pinto, Y. Dan, Cell-Type-Specific Activity in Prefrontal Cortex during Goal-Directed Behavior. Neuron 87, 437–450 (2015).

14. J. M. Otis, V. M. Namboodiri, A. M. Matan, E. S. Voets, E. P. Mohorn, O. Kosyk, J. A. McHenry, J. E. Robinson, S. L. Resendez, M. A. Rossi, G. D. Stuber, Prefrontal cortex output circuits guide reward seeking through divergent cue encoding. Nature 543, 103–107 (2017).

15. W. E. Pratt, S. J. Y. Mizumori, Neurons in rat medial prefrontal cortex show anticipatory rate changes to predictable differential rewards in a spatial memory task. Behavioural Brain Research 123, 165–183 (2001).

16. A. Pasupathy, E. K. Miller, Different time courses of learning-related activity in the prefrontal cortex and striatum. Nature 433, 873–876 (2005).

17. M. J. Wojcik, J. P. Stroud, D. Wasmuht, M. Kusunoki, M. Kadeisha, M. J. Buckley, R. P. Costa, N. E. Myers, L. T. Hunt, J. Duncan, M. G. Stokes, Learning shapes neural geometry in the primate prefrontal cortex. Nat Neurosci 10.1038/s41593-026-02333-w (2026).

18. L. N. Driscoll, N. L. Pettit, M. Minderer, S. N. Chettih, C. D. Harvey, Dynamic Reorganization of Neuronal Activity Patterns in Parietal Cortex. Cell 170, 986–999 e916 (2017).

19. M. E. Rule, A. R. Loback, D. V. Raman, L. N. Driscoll, C. D. Harvey, T. O’Leary, Stable task information from an unstable neural population. Elife 9 (2020).

20. J. Taxidis, E. A. Pnevmatikakis, C. C. Dorian, A. L. Mylavarapu, J. S. Arora, K. D. Samadian, E. A. Hoffberg, P. Golshani, Differential Emergence and Stability of Sensory and Temporal Representations in Context-Specific Hippocampal Sequences. Neuron 108, 984–998 e989 (2020).

21. J. R. Climer, H. Davoudi, J. Y. Oh, D. A. Dombeck, Hippocampal representations drift in stable multisensory environments. Nature 10.1038/s41586-025-09245-y (2025).

22. W. Mau, D. W. Sullivan, N. R. Kinsky, M. E. Hasselmo, M. W. Howard, H. Eichenbaum, The Same Hippocampal CA1 Population Simultaneously Codes Temporal Information over Multiple Timescales. Curr Biol 28, 1499–1508 e1494 (2018).

23. C. Clopath, T. Bonhoeffer, M. Hubener, T. Rose, Variance and invariance of neuronal long-term representations. Philos Trans R Soc Lond B Biol Sci 372 (2017).

24. P. D. Rich, S. Y. Thiberge, N. D. Daw, D. W. Tank, Error-driven changes in hippocampal representations accompany flexible re-learning. *bioRxiv* 10.1101/2025.05.20.655046 (2025).

25. L. N. Driscoll, L. Duncker, C. D. Harvey, Representational drift: Emerging theories for continual learning and experimental future directions. Curr Opin Neurobiol 76, 102609 (2022).

26. M. Natrajan, J. E. Fitzgerald, Stability through plasticity: Finding robust memories through representational drift. Proc Natl Acad Sci U S A 122, e2500077122 (2025).

27. Y. Li, W. Yin, X. Wang, J. Li, S. Zhou, C. Ma, P. Yuan, B. Li, Stable sequential dynamics in prefrontal cortex represents subjective estimation of time. eLife 13, RP96603 (2024).

28. H. Muysers, H. L. Chen, J. Hahn, S. Folschweiller, T. Sigurdsson, J. F. Sauer, M. Bartos, A persistent prefrontal reference frame across time and task rules. Nat Commun 15, 2115 (2024).

29. O. C. Sylte, H. Muysers, H. L. Chen, M. Bartos, J. F. Sauer, Neuronal tuning to threat exposure remains stable in the mouse prefrontal cortex over multiple days. PLoS Biol 22, e3002475 (2024).

30. C. K. Machens, R. Romo, C. D. Brody, Functional, but not anatomical, separation of “what” and “when” in prefrontal cortex. J Neurosci 30, 350–360 (2010).

31. T. W. Bernklau, B. Righetti, L. S. Mehrke, S. N. Jacob, Striatal dopamine signals reflect perceived cue-action-outcome associations in mice. Nat Neurosci 27, 747–757 (2024).

32. E. A. Pnevmatikakis, D. Soudry, Y. Gao, T. A. Machado, J. Merel, D. Pfau, T. Reardon, Y. Mu, C. Lacefield, W. Yang, M. Ahrens, R. Bruno, T. M. Jessell, D. S. Peterka, R. Yuste, L. Paninski, Simultaneous Denoising, Deconvolution, and Demixing of Calcium Imaging Data. Neuron 89, 285–299 (2016).

33. A. Giovannucci, J. Friedrich, P. Gunn, J. Kalfon, B. L. Brown, S. A. Koay, J. Taxidis, F. Najafi, J. L. Gauthier, P. Zhou, B. S. Khakh, D. W. Tank, D. B. Chklovskii, E. A. Pnevmatikakis, CaImAn an open source tool for scalable calcium imaging data analysis. Elife 8 (2019).

34. M. G. Stokes, M. Kusunoki, N. Sigala, H. Nili, D. Gaffan, J. Duncan, Dynamic coding for cognitive control in prefrontal cortex. Neuron 78, 364–375 (2013).

35. A. Bellafard, G. Namvar, J. C. Kao, A. Vaziri, P. Golshani, Volatile working memory representations crystallize with practice. Nature 629, 1109–1117 (2024).

36. X.-X. Lin, A. Nieder, S. N. Jacob, The neuronal implementation of representational geometry in primate prefrontal cortex. Science Advances 9 (2023).

37. A. H. Williams, T. H. Kim, F. Wang, S. Vyas, S. I. Ryu, K. V. Shenoy, M. Schnitzer, T. G. Kolda, S. Ganguli, Unsupervised Discovery of Demixed, Low-Dimensional Neural Dynamics across Multiple Timescales through Tensor Component Analysis. Neuron 98, 1099–1115 e1098 (2018).

38. D. Kobak, W. Brendel, C. Constantinidis, C. E. Feierstein, A. Kepecs, Z. F. Mainen, X. L. Qi, R. Romo, N. Uchida, C. K. Machens, Demixed principal component analysis of neural population data. Elife 5 (2016).

39. R. V. Rikhye, A. Gilra, M. M. Halassa, Thalamic regulation of switching between cortical representations enables cognitive flexibility. Nat Neurosci 21, 1753–1763 (2018).

40. L. I. Schmitt, R. D. Wimmer, M. Nakajima, M. Happ, S. Mofakham, M. M. Halassa, Thalamic amplification of cortical connectivity sustains attentional control. Nature 545, 219–223 (2017).

41. J. M. Birrell, V. J. Brown, Medial Frontal Cortex Mediates Perceptual Attentional Set Shifting in the Rat. The Journal of Neuroscience 20, 4320–4324 (2000).

42. R. Dias, T. W. Robbins, A. C. Roberts, Dissociation in prefrontal cortex of affective and attentional shifts. Nature 380, 69–72 (1996).

43. A. Izquierdo, Functional Heterogeneity within Rat Orbitofrontal Cortex in Reward Learning and Decision Making. J Neurosci 37, 10529–10540 (2017).

44. A. Pellegrino, H. Stein, N. A. Cayco-Gajic, Dimensionality reduction beyond neural subspaces with slice tensor component analysis. Nat Neurosci 27, 1199–1210 (2024).

45. M. C. Aoi, V. Mante, J. W. Pillow, Prefrontal cortex exhibits multidimensional dynamic encoding during decision-making. Nat Neurosci 23, 1410–1420 (2020).

46. V. Mante, D. Sussillo, K. V. Shenoy, W. T. Newsome, Context-dependent computation by recurrent dynamics in prefrontal cortex. Nature 503, 78–84 (2013).

47. A. Dubreuil, A. Valente, M. Beiran, F. Mastrogiuseppe, S. Ostojic, The role of population structure in computations through neural dynamics. Nat Neurosci 25, 783–794 (2022).

48. J. Soldado--Magraner, V. Mante, M. Sahani, Inferring context--dependent computations through linear approximations of prefrontal cortex dynamics. Sci. Adv. 10, eadl4743 (2024).

49. M. Rigotti, O. Barak, M. R. Warden, X. J. Wang, N. D. Daw, E. K. Miller, S. Fusi, The importance of mixed selectivity in complex cognitive tasks. Nature 497, 585–590 (2013).

50. O. Barak, M. Rigotti, S. Fusi, The sparseness of mixed selectivity neurons controls the generalization-discrimination trade-off. J Neurosci 33, 3844–3856 (2013).

51. S. Bernardi, M. K. Benna, M. Rigotti, J. Munuera, S. Fusi, C. D. Salzman, The Geometry of Abstraction in the Hippocampus and Prefrontal Cortex. Cell 183, 954–967 e921 (2020).

52. M. Saez, R. Blassberg, E. Camacho-Aguilar, E. D. Siggia, D. A. Rand, J. Briscoe, Statistically derived geometrical landscapes capture principles of decision-making dynamics during cell fate transitions. Cell Syst 13, 12–28 e13 (2022).

53. J. X. Wang, Z. Kurth-Nelson, D. Kumaran, D. Tirumala, H. Soyer, J. Z. Leibo, D. Hassabis, M. Botvinick, Prefrontal cortex as a meta-reinforcement learning system. Nat Neurosci 21, 860–868 (2018).

54. W. Schultz, P. Dayan, R. P. Montague, A Neural Substrate of Prediction and Reward. Science 275 (1997).

55. P. P. Witkowski, S. A. Park, E. D. Boorman, Neural mechanisms of credit assignment for inferred relationships in a structured world. Neuron 110, 2680–2690 e2689 (2022).

56. M. L. Mack, A. R. Preston, B. C. Love, Ventromedial prefrontal cortex compression during concept learning. Nat Commun 11, 46 (2020).

57. J. Hirokawa, A. Vaughan, P. Masset, T. Ott, A. Kepecs, Frontal cortex neuron types categorically encode single decision variables. Nature 576, 446–451 (2019).

58. M. Wilhelm, Y. Sych, A. Fomins, J. L. Alatorre Warren, C. Lewis, L. Serratosa Capdevila, R. Boehringer, E. A. Amadei, B. Grewe, E. C. O’Connor, B. J. Hall, F. Helmchen, Striatum-projecting prefrontal cortex neurons support working memory maintenance. Nat Commun 14, 7016 (2023).

59. S. L. Jackman, C. H. Chen, S. N. Chettih, S. Q. Neufeld, I. R. Drew, C. K. Agba, I. Flaquer, A. N. Stefano, T. J. Kennedy, J. E. Belinsky, K. Roberston, C. C. Beron, B. L. Sabatini, C. D. Harvey, W. G. Regehr, Silk Fibroin Films Facilitate Single-Step Targeted Expression of Optogenetic Proteins. Cell Rep 22, 3351–3361 (2018).

60. K. B. Franklin, G. Paxinos, Paxinos and Franklin’s the Mouse brain in stereotaxic coordinates, compact: The coronal plates and diagrams. (Academic press, 2019).

